# Anisotropic Crb accumulation, modulated by Src42A, orients epithelial tube growth in *Drosophila*

**DOI:** 10.1101/287854

**Authors:** Ivette Olivares-Castiñeira, Marta Llimargas

## Abstract

Size control of internal tubular organs, such as the lungs or vascular system, is critical for proper physiological activity and to prevent disease or malformations. This control incorporates the intrinsic physical anisotropy of tubes to generate proportionate organs that match their function. The exact mechanisms underlying tube size control and how tubular anisotropy is translated at the cellular level are still not fully understood. Here we investigate these mechanisms using the *Drosophila* tracheal system. We show that the apical polarity protein Crumbs transiently accumulates anisotropically at longitudinal cell junctions during tube elongation. We provide evidence indicating that the accumulation of Crumbs in specific apical domains correlates with apical surface growth. Hence, we propose that the anisotropic accumulation of Crb at the cellular level promotes a bias towards longitudinal membrane growth, orienting cell elongation and, as a consequence, longitudinal growth at the tissue level. We find that Src42A promotes Crb anisotropic accumulation, thereby identifying the first polarised cell behaviour downstream of Src42A. Our results indicate that Src42A activity favours a higher turnover of Crb protein at longitudinal junctions, possibly through a mechanism regulating protein trafficking and/or recycling. We propose a model where Src42A would sense the inherent anisotropic mechanical tension of the tube and translate it into a polarised Crumbs-mediated membrane growth, orienting longitudinal tube growth. This work provides new insights into the key question of how organ growth is controlled and polarised and unveils the function of two conserved proteins, Crumbs and Src42A, with important roles in development and homeostasis as well as in disease, in this biological process.

## Introduction

Tubes are physically anisotropic, with a curved circumferential axis and a flat longitudinal one. This physical property confers orientation and polarisation to tubes, critical for their physiological activity in biological systems. Many vital organs, such as the lungs, vascular system or mammary glands, are internal tubular structures (1–3), underscoring the importance of investigating how they form and polarise to be functional.

The tracheal (respiratory) system of *Drosophila* is a paradigm for the analysis of tubular organs (4). After a morphogenetic phase by branching morphogenesis (5, 6) the tracheal tubes mature and become physiologically active (7). Tube maturation ensures the acquisition of the correct tube diameter and length, and of gas filling, critical steps for organ functional activity (3, 8, 9). The diametrical and longitudinal growth of tracheal tubes are two different events regulated by different mechanisms (reviewed in (10)). Tube longitudinal growth starts at the end of stage 14 and continues until the embryo hatches. Several mechanisms have been shown to control this elongation. These include the proper modification of the apical extracellular matrix, aECM, consisting of a transient filament made of chitin and chitin-associated proteins (11, 12)), particularly Serpentine (Serp) and Vermiform (Verm), whose absence leads to tube overelongation (13, 14). Cell intrinsic mechanisms such as Crumbs (Crb)-mediated apical membrane growth (15, 16) and Src42A-mediated cell shape regulation (17, 18) also control tube length. A model of tube length control has been proposed (10, 15, 19) where the apical membrane expansion force driven by Crb is balanced with the aECM resistance through factors that attach the aECM to the apical membrane. Excess or defects in any of these two forces or their uncoupling leads to tube length defects. However, several questions remain unclear, such as how the Crb-mediated apical membrane growth is biased to the longitudinal direction, how the different factors interact, or how uniform Src42A accumulation controls polarised cell shape changes.

## Results and Discussion

### Crb accumulates anisotropically during tube elongation

Since Crb has been proposed to regulate tube length by promoting apical membrane growth (15, 16), we first examined Crb accumulation in the Dorsal Trunk (DT, the main tracheal trunk connecting to the exterior through the spiracles). Crb can localise to different subdomains of the apical membrane during tracheal development: the SubApical Region (SAR) and the Apical Free Region (AFR) (20). The AFR corresponds to the most apical domain, free of contact with other epithelial cells and in direct contact with the lumen in the case of tubular organs like the trachea, while the SAR corresponds to the most apicolateral membrane domain of contact between neighboring epithelial cells. We previously showed that during the stages of higher longitudinal DT growth, stage 15 onwards, Crb accumulated strongly in the SAR, displaying a mesh-like pattern that identifies the apical junctional domain (20). Strikingly, we now observed that Crb was anisotropically (not uniformly) distributed in the SAR of cell junctions. We classified cell junctions as longitudinal cell junctions (LCJs), mainly parallel to the longitudinal axis of the tube, and transverse cell junctions (TCJs), perpendicular to the longitudinal axis (see Materials and Methods). We found that Crb accumulation was more visible at LCJs than at TCJs (Fig 1A, S1A,B), observing several examples where accumulation of Crb at TCJs was almost absent (Fig 1A yellow arrowheads). We quantified the accumulation of Crb (total pixels/junctional length) at LCJs and TCJs and found that Crb accumulation was clearly biased to LCJs, where levels were around 30% higher than at TCJs (average % of difference of Crb accumulation at LCJs and TCJs, n=15 embryos). To compare different embryos we calculated the LCJ/TCJ ratio of Crb accumulation (Fig. 1B), which showed an average of 1,5, indicating that Crb is anisotropically distributed, i.e. planar polarised. In contrast to Crb, *D*E-Cadherin (*D*E-cad), a core component of the Adherens Junctions (AJs), was equally distributed among all cell junctions (Fig. 1A, S1A,B). The ratio of accumulation in LCJ/TCJ was close to 1 (Fig 1B), indicating that the anisotropic distribution is not a general feature of all apical junctional proteins. These results indicated that a larger proportion of LCJs accumulate higher levels of Crb than TCJs. Interestingly, this transient planar polarisation of Crb accumulation observed at stages 15-16 correlated with the anisotropic growth of the tracheal tubes along the longitudinal axis of the DT. Because Crb has been proposed to control tube growth through apical membrane expansion (15, 16), our results raise the possibility that a planar polarised Crb accumulation drives the polarised longitudinal tube growth. It is worth pointing out that anisotropies of Crb, like the one described here, or of other apical determinants, have important implications in morphogenesis (21, 22).

**Figure 1:**
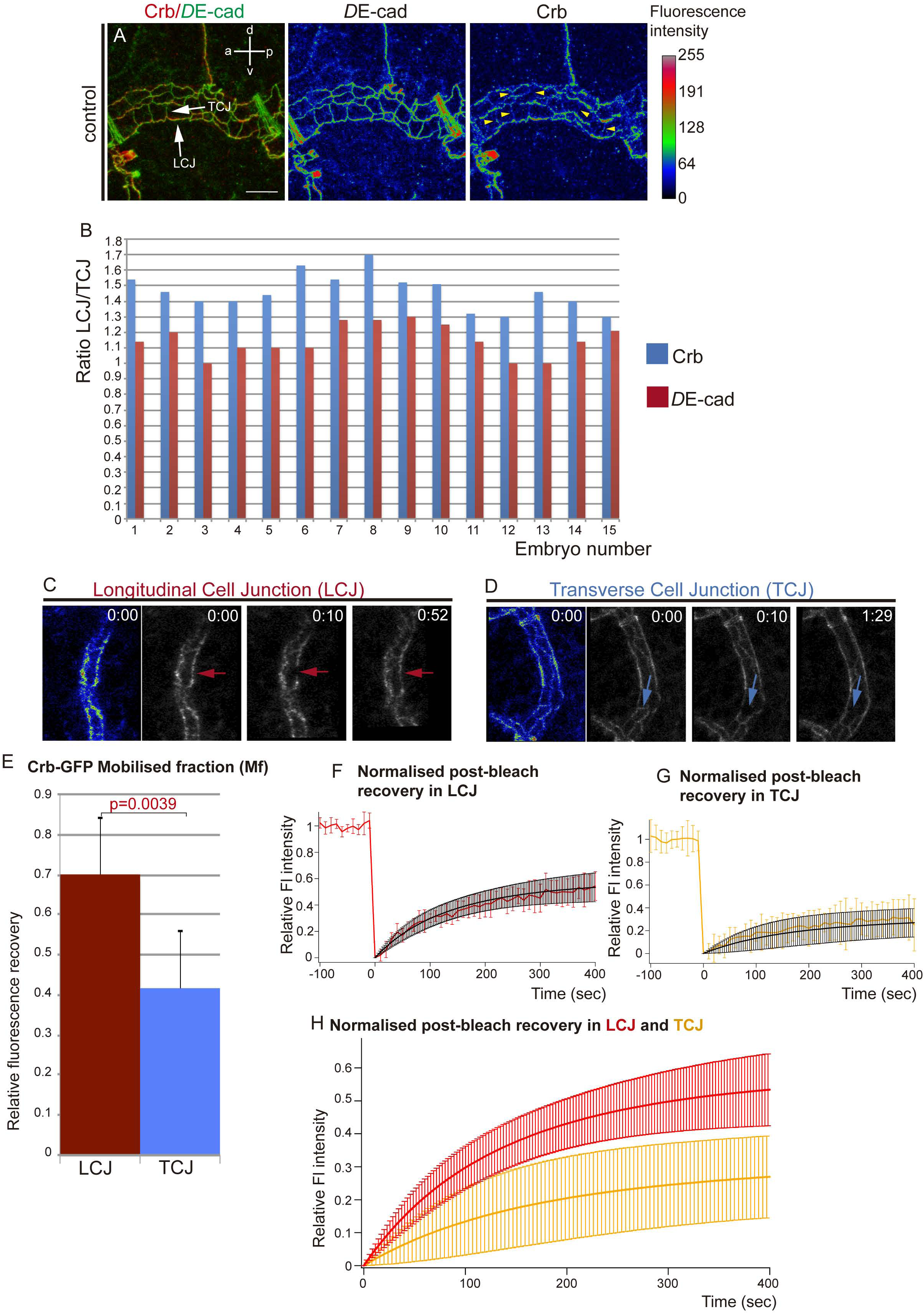
Anisotropic accumulation of Crb during tube elongation. (A) Confocal projection of DT cells of stage 16 embryos stained for *D*E-cad and Crb. The fluorescence intensity of *D*E-cad and Crb is shown in heat maps. The fluorescence intensity values mapped to the gradient bar to the right ranging from black (0) to white (255). The gradient bar was generated with the “calibration bar” tool in ImageJ. Note that while *D*E-cad shows similar fluorescence intensity at longitudinal and transverse cell junctions (LCJs and TCJs respectively, white arrows), Crb fluorescence intensity is higher at LCJs. Crb accumulation at many TCJs is hardly detectable (yellow arrowheads). Scale bar 7,5 μm (B) Quantification of the ratio of *D*E-cad and Crb accumulation at LCJs and TCJs for each embryo analysed. Note that while *D*E-cad ratio is close to 1, indicating an equal fluorescence intensity at all cell junctions, Crb ratio indicates increased fluorescence intensity in LCJs. n=254 LCJs and n=314 TCJs from 15 wild type embryos. (C-H) FRAP experiments of Crb-GFP (a viable knock-in allele that provides the only source of Crb protein in the embryo) at LCJs or TCJs in the DT of stage 16 embryos. (C,D) Images acquired before (time 0 min) and after (from time 10 min) photobleaching a region in an LCJ (red arrows in D) or TCJ (blue arrows in E) are shown. The initial fluorescence intensity is shown in heat maps at the left. Note the difference in fluorescence recovery in LCJs and TCJs. (E) The Mobile fraction, Mf, was calculated and compared for LCJs and TCJs experiments. Mf was significantly higher at LCJs than at TCJs. Error bars indicate standard error (s.e.). (F,G) The levels of fluorescence relative to the pre-bleach levels (from -100 to 0 sec) during the FRAP experiments (from 0 to 400 sec) are shown for each experimental situation as the average of the different experiments (in red for LCJs in G, and orange for TCJs in H). The values obtained were fit to a single exponential curve (in black), indicating one single pool of Crb-GFP recovery. (H) Comparison of the fit of normalised post-bleach recovery in LCJs and TCJs. Note the higher recovery at LCJs. Error bars in G-I indicate standard deviation (s.d.). n=8 LCJs and n=6 TCJs from wild type embryos.

### Differential turnover of Crb protein at cell junctions

Different molecular mechanisms could underlie the preferential accumulation of Crb at LCJs, such as specific Crb degradation at TCJs, specific stabilisation at LCJs, targeted intracellular trafficking, differential protein recycling, among others. To investigate the dynamics of Crb protein we performed FRAP analysis at either LCJs or TCJs of embryos carrying a viable and functional *CrbGFP* allele (Fig. 1C,D, Movies 1,2). We found that the amount of fluorescent protein, relative to the pre-bleach value, mobilized during the experimental time (mobile fraction, Mf) was significantly higher at LCJs compared to TCJs (Fig. 1E), indicating a higher recovery of Crb-GFP protein at LCJs (Fig. 1E-H, S2D,E). To assess the recovery kinetics we calculated the half-time (t_1/2_, time to reach half of the Mf). We found that the half-time was not significantly different at LCJs and TCJs, suggesting that the recovery rate is comparable at the differently oriented junctions (Fig. S2B). Moreover, kymographs of the bleached regions suggested that the recovery is not due to lateral diffusion (Fig. S2F,G). Altogether our results are consistent with a model where the turnover of Crb protein is higher at LCJs. As we had previously described an endocytic-recycling turnover of Crb operating during tracheal development (20), the results suggest that Crb would be more prone to recycle and/or traffic to/at LCJs than to/at TCJs.

### Src42A promotes Crb anisotropic accumulation

We next asked how the anisotropic distribution of Crb is regulated. To investigate this question we turned our attention to Src42A, as it triggers one of the mechanisms regulating tube elongation, orienting membrane growth on the longitudinal axis. In conditions of Src42A loss of function, LCJs do not expand and tubes become shorter (17, 18). We analysed Crb accumulation in embryos expressing a dominant negative form of Src42A (Src42^DN^) in the trachea and observed a more uniform distribution at LCJs and TCJs (Fig. 2A, S1C,D). Quantification of Crb levels indicated that while there was still some increased accumulation of Crb at LCJs compared to TCJs, the differences were reduced. Analysis of the LCJ/TCJ ratio of Crb clearly showed a significant decrease when compared to the control (Fig 2B,C), indicating a more uniform accumulation in Src42A^DN^ conditions. *D*E-cad accumulation remained unchanged in Src42^DN^ as compared to the control (Fig. 2A-C). Altogether these results show that a decrease in Src42A activity leads to a decrease of the anisotropic accumulation of Crb.

**Figure 2:**
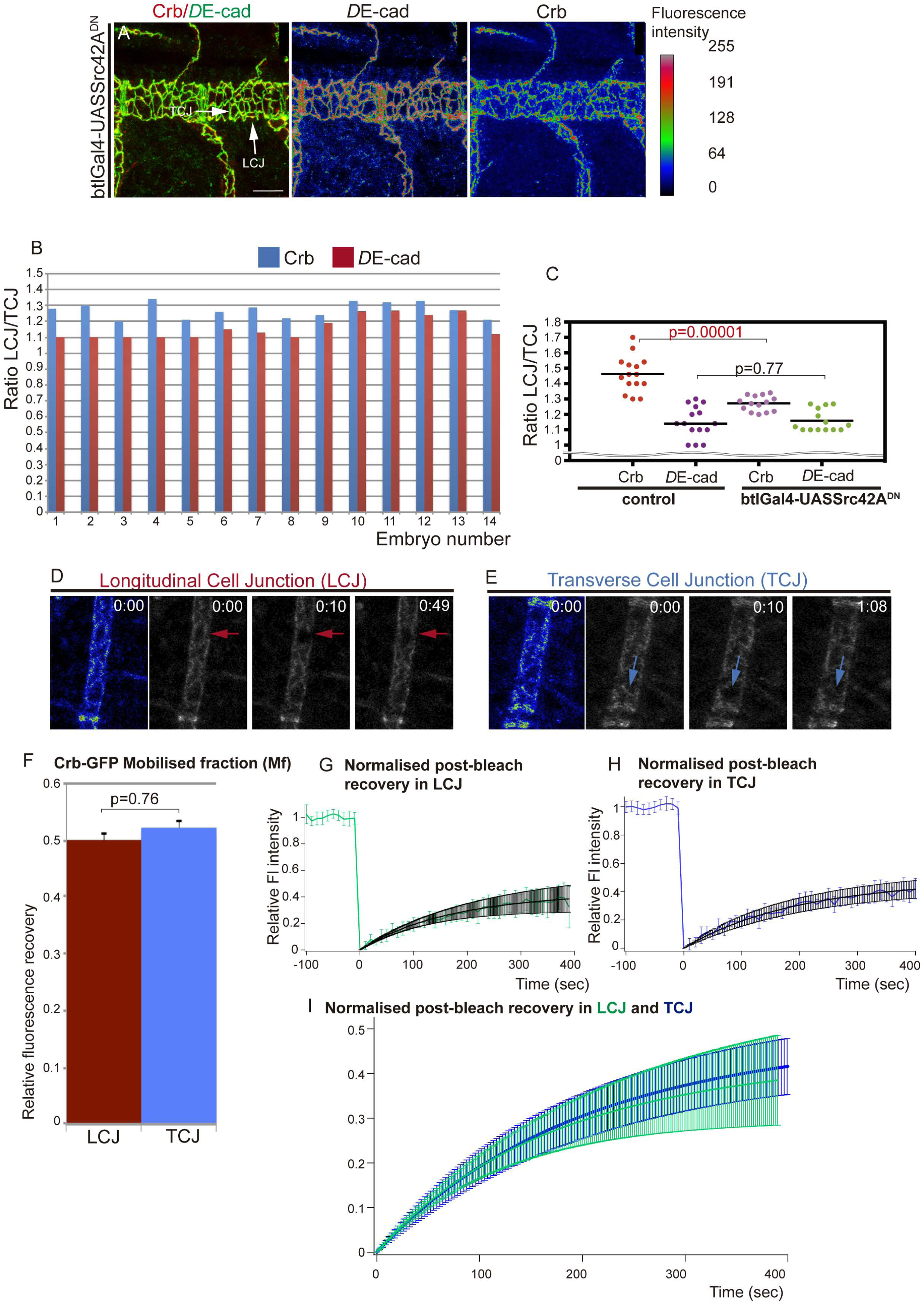
Src42A modulates Crb anisotropic accumulation. (A) Confocal projection of DT cells of stage 16 embryos in which Src42A is downregulated in the tracheal cells. Embryos are stained for *D*E-cad and Crb. The colour-coded fluorescence intensity shown for *D*E-cad and Crb matches the heat map shown on the right. Note the homogeneity of fluorescence intensities at LCJs and TCJs for *D*E-cad and also for Crb. Scale bar 10 μm (B) Quantification of the ratio of *D*E-cad and Crb accumulation at LCJs and TCJs in each embryo analysed. n=349 LCJs and n=417 TCJs from 14 mutant embryos. (C) Scatter Plot comparing the LCJ/TCJ ratio of accumulation of Crb and *D*E-cad in control and Src^DN^ embryos. Note that while the ratio of *D*E-cad is comparable in control and mutants, the ratio of LCJ/TCJ Crb accumulation is significantly different in the two conditions. (D-I) FRAP experiments of Crb-GFP in LCJs or TCJs in the DT of stage 16 embryos in which Src42A is downregulated in the tracheal cells. (D,E) Images acquired before (time 0 min) and after (from time 10 min) photobleaching a region in a LCJ (red arrows in D) or a TCJ (blue arrows in E) are shown. The initial fluorescence intensity is shown in heat maps at the left. Note the comparable fluorescence recovery in LCJs and TCJs. (F) The Mobile fraction in LCJs and TCJs was comparable. Error bars indicate standard error (s.e.). (G,H) The levels of fluorescence relative to the pre-bleach levels during the FRAP experiments are shown for each experimental situation as the average of the different experiments (in green for LCJs in G, and blue for TCJs in H). The values obtained were fit to a single exponential curve (in black). (I) Comparison of the fit of normalised post-bleach recovery in LCJs and TCJs. Note the similar recovery in LCJs and TCJs. Error bars in G-I indicate standard deviation (s.d.). n=7 LCJs and n=9 TCJs from mutant embryos.

To further explore Src42A requirement we performed FRAP experiments in *Crb-GFP* embryos in which Src42A was downregulated (Fig. 2D,E, Movies 3,4). We found clear differences with respect to control: while in the control the Mf and recovery curves of LCJs and TCJs were clearly different (Fig. 1E-H, S2C), in Src42^DN^ conditions the Mf and recovery curves of LCJs and TCJs were comparable (Fig. 2F-I, S2C,H,I). The Mf at the LCJs of Src42^DN^ was significantly lower than the Mf at the LCJs in control embryos, and was similar to the Mf at TCJs in control and mutant embryos (Fig. S2A). The fact that Crb recovery is affected particularly at LCJs when Src42A is downregulated strongly suggests that Src42A is (more) active precisely at LCJs, as previously suggested (17, 18). The halftime recovery, t_1/2_, was comparable to that of control embryos, indicating a recovery rate similar in all cases (Fig. S2B). Again, kymographs suggested no major contribution of lateral diffusion to fluorescence recovery (Fig. S2J,K).

Altogether our results indicate that Src42A contributes to Crb preferential accumulation at LCJs. Src42A controls tube elongation through interactions with dDaam and the remodelling of AJs (17, 18). Here we propose that Crb would also act downstream of Src42A contributing to this function. Interestingly, we find that Src42A does not regulate the base level accumulation of Crb in the SAR or its kinetics, but instead it promotes Crb enrichment at LCJs. Thus, a Src42A-independent accumulation of Crb in tracheal cells together with other Src42A-independent mechanisms of apical membrane growth may be responsible for tube growth in the absence of Src42A.

### Regulation of tube elongation by constitutive activation of Src42A

We found that Src42A promotes Crb accumulation at LCJs. We then asked whether the overelongation of tubes observed in Src42A overactivation (either overexpression or constitutive activation) (17, 18) was due to an increased accumulation of Crb at LCJs. In contrast to this expectation we found that Crb was strongly decreased in the SAR of DT cells (Fig. 3A, B). This result suggested that the tube length defects produced by Src42A overactivation were caused by a mechanism different than the one operating in physiological conditions. To investigate this further we analysed other known tube length regulators. One of them is Serp, which regulates the aECM organisation (13, 14). We found that in Src42A^CA^ conditions, Serp is completely lost from the luminal compartment

**Figure 3:**
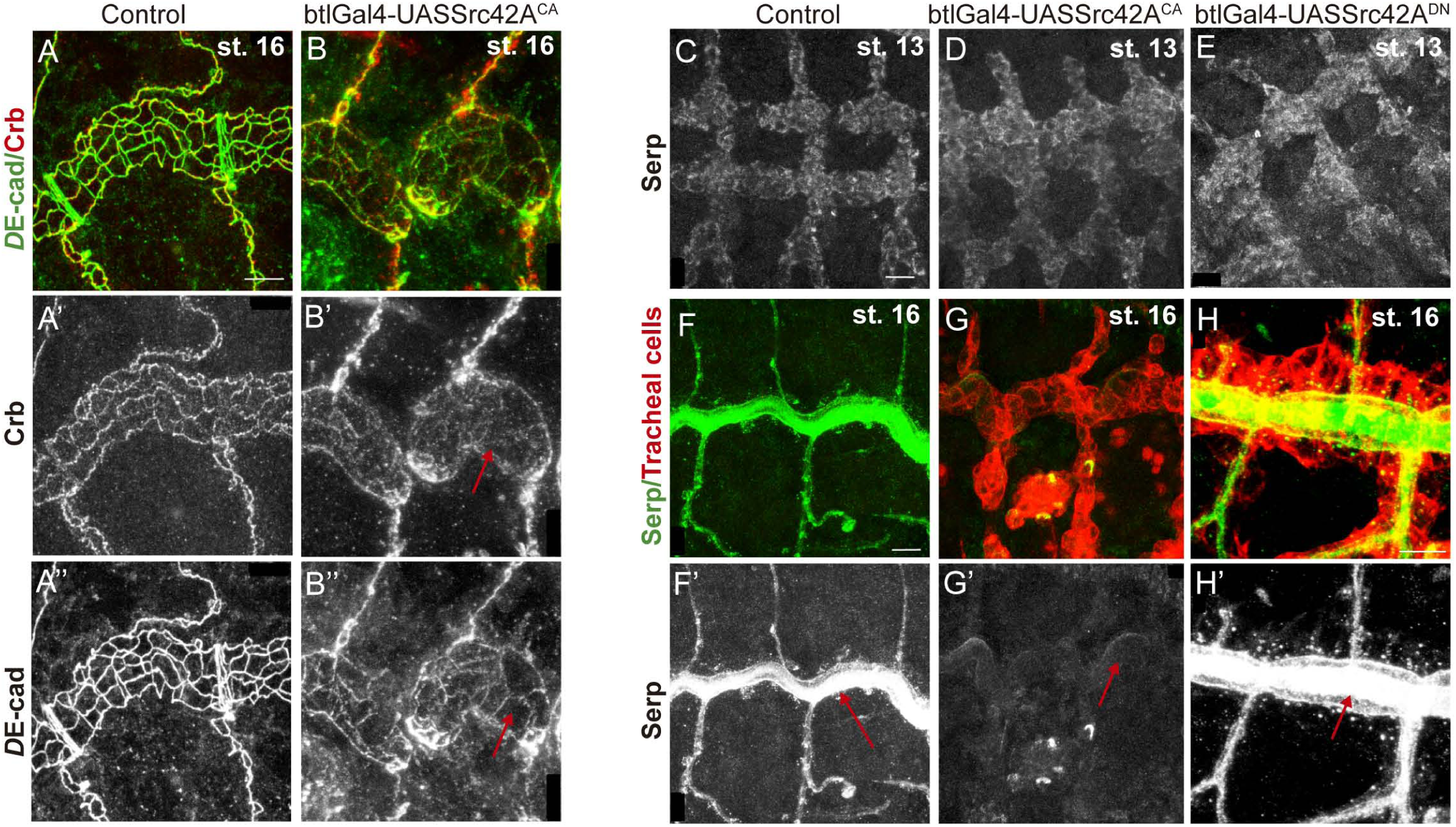
Src42A overactivation affects the accumulation of proteins that undergo endocytic trafficking in the tracheal system. Lateral views showing a region of the DT of embryos of indicated genotypes stained for the indicated markers. Note that Crb accumulation is decreased in the DT (red arrow in B’) and *D*E-cad staining is fragmented and decreased in AJs (red arrow in B’’) in embryos expressing constitutively active Src42A. Serp is accumulated normally in tracheal cells at early stages (C,D), but it is later lost from the luminal compartment (red arrows in E,F) in Src42A^CA^ embryos. In contrast, Serp accumulation is not affected when Src42A is downregulated (E,H) and it is normally detected in the lumen at late stages (red arrow in H’). Scale bar A,H 7,5 μm. C,F 10 μm

(Fig. 3F,G), although, as in wild type, tracheal cells accumulate Serp at early stages (Fig 3C, D). This result provides explanation for the tube elongation defects observed in Src42A^CA^ conditions, as Serp absence leads to tube overelongation. Interestingly, we could not detect defects in Serp accumulation in Src42A^DN^ conditions (Fig 3E, H), as previously reported (18). These results indicate that Src42A overactivation use a different mechanism than the one used in physiological conditions to drive tube elongation. These observations support the notion that tube elongation depends on the activity and coordination of different mechanisms (19), and indicate that Src is able to regulate several of them.

These results also pointed to a possible role of Src42A regulating protein trafficking, as the tracheal accumulation of several proteins regulated by Src42A, such as Crb and Serp (Fig. 2, 3B’,G’) and *D*E-cad (Fig 3B’’ and (17)), have been shown to depend on their intracellular trafficking (20, 23, 24). Roles for Src42A in protein trafficking have been proposed in different contexts (17, 25–27). Src42A could regulate protein trafficking directly, or indirectly through the regulation of the actin cytoskeleton. The actin cytoskeleton plays a capital role in protein trafficking (28) and Src42A acts as a regulator of the actin cytoskeleton (29, 30). A disruption of actin organisation in Src42A^CA^ could lead to defects in the sorting of different cargoes as well as defects in endosomal maturation.

### Crb accumulation in the SAR correlates with apical cell expansion

Independently of our finding of Crb anisotropic distribution and its regulation by Src42A, we also aimed to investigate a possible role of Crb in cell elongation. Crb has been proposed to promote apical membrane growth independently of its role in apicobasal polarity at late stages of epithelial differentiation (31, 32), but the exact mechanism is not fully understood. We further analysed this role by investigating the elongation of the tracheal cells during tube elongation. As above-mentioned, during tracheal development Crb becomes enriched in the SAR during the stages of DT elongation (20). This observation raises the possibility that Crb accumulation in the SAR drives, favours or facilitates apical membrane growth. To investigate this hypothesis we analysed Crb accumulation in the SAR or AFR in conditions in which apicobasal polarity was unaffected. In the first set of experiments we used the tracheal system. We had previously shown that in tracheal cells EGFR plays a role in regulating Crb subcellular accumulation and that the tracheal expression of a constitutively active form of EGFR, EGFR^CA^, leads to a loss of Crb in the SAR and a concomitant increased accumulation in the AFR (Fig. 4A, B, S1E’-F’). Quantification of the ratio of Crb accumulation in the SAR versus the AFR in control and EGFR^CA^ expressing tracheal cells clearly indicated a significant decrease in the mutant condition (Fig. 4C) (20). We now extended this analysis and quantified the apical surface area. This quantification also indicated a clear difference (Fig. 4D), with cells expressing EGFR^CA^ showing a smaller apical area than control ones (Fig. 4A’,B’, S1E-F). In a second set of experiments we used another tubular epithelial tissue, the salivary gland (SG), to investigate whether Crb subcellular accumulation and apical expansion were also correlated. We found that in SGs of control embryos, and in contrast to the tracheal tissue, Crb was high in the AFR, with little accumulation in the SAR (Fig. 4E,G, S1G’,H’). Because in the trachea EGFR plays a role in Crb subcellular accumulation, we explored if it also controlled Crb accumulation in the SG. Interestingly, when we blocked EGFR activity in the SG, by expressing EGFR^DN^, we found a clear relocalisation and accumulation of Crb in the SAR (Fig. 4F,G S1I’,J’). SG cells expressing EGFR^DN^ showed increased apical surface area (Fig. 4F’,H, S1I,J), which was accompanied by abnormal SG morphology. The results indicated that manipulation of EGFR activity has an effect on apical surface area. We showed that EGFR regulates the trafficking of different cargoes, in particular Crb and Serp (20), raising the possibility that the regulation of the apical surface area depends on targets different than Crb. However, the fact that Serp is not present in the SGs and that Crb has already been proposed to promote apical membrane growth (31, 32), strongly suggest that Crb is at least one of the targets downstream of EGFR regulating apical growth.

**Figure 4:**
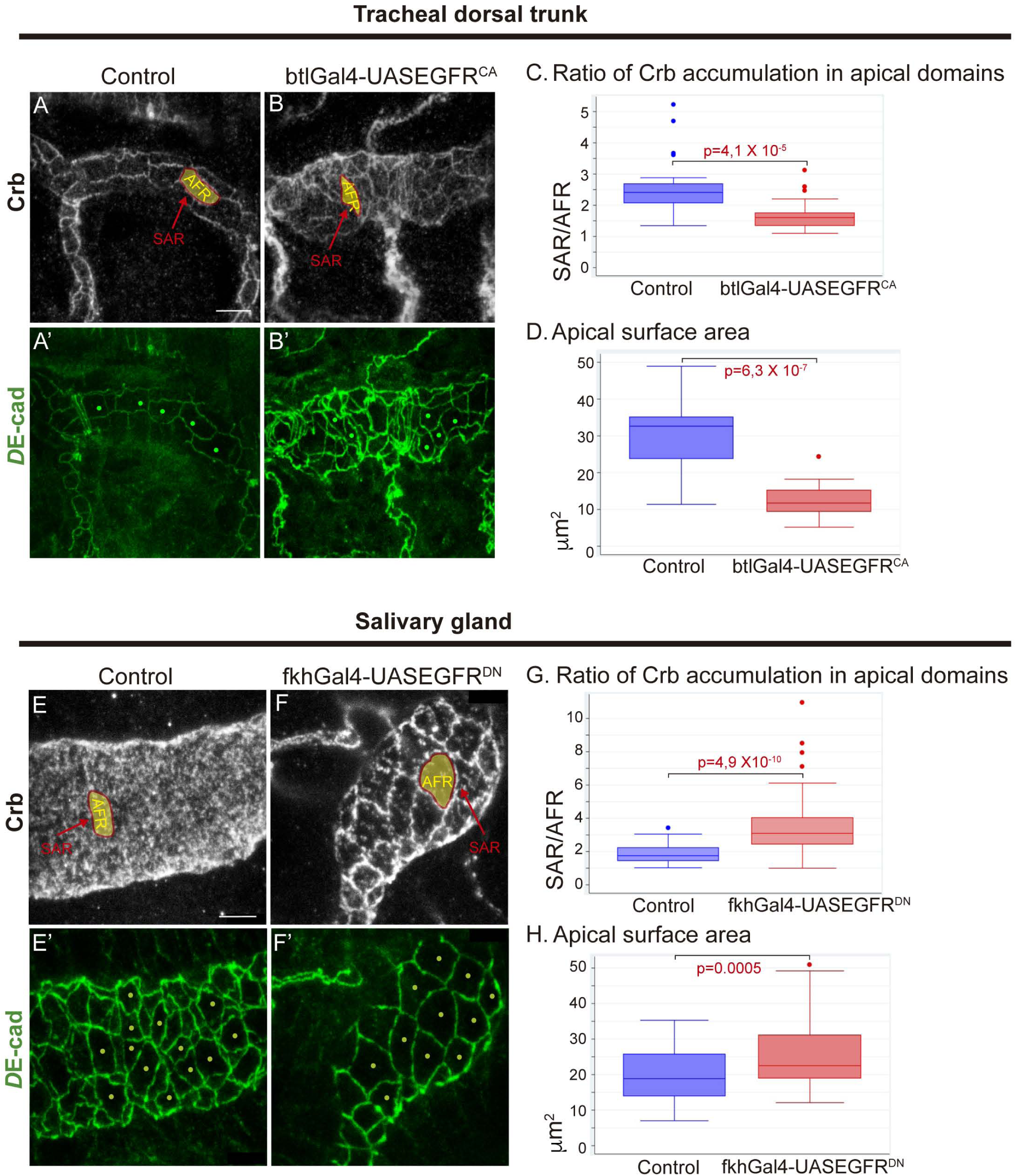
The subcellular localisation of Crb in apical domains correlate with apical expansion. (A-B) Lateral views showing a region of the DT of stage 16 embryos of indicated genotypes stained for the indicated markers. The SAR outlines the cell contours and the AFR fills the cell space inside the SAR. Note the poor enrichment of Crb in the SAR when EGFR is constitutively activated. *D*E-cad stains the junctional apical area homogeneously, allowing the measurement of the apical surface area in selected cells (little green dots). Scale bar 7,5 μm (C) Quantification of the ratio of Crb accumulation in the different apical regions (as published in (20)). n=25 cells from 8 control embryos, n=53 cells from 7 EGFR^CA^ embryos (D) Quantification of the area of the apical region. n=17 cells from 4 control embryos, n=25 cells from 5 EGFR^CA^ embryos (E-F) Lateral views showing a region of the salivary gland of stage 16 embryos of indicated genotypes stained for the indicated markers. Note that in control embryos Crb is not particularly enriched in the SAR (E). In contrast, when EGFR is downregulated (F) Crb clearly relocalises and accumulates in the SAR and the apical domain of the cells (green dots) is enlarged. Scale bar 5 μm (G) Quantification of the ratio of Crb accumulation in the different apical regions. n=30 cells from 7 control embryos, n=67 cells from 9 EGFR^DN^ embryos (H) Quantification of the area of the apical region. n=54 cells from 7 control embryos, n=69 cells from 9 EGFR^DN^ embryos

Altogether these results confirm a role of EGFR in regulating the accumulation of Crb in the SAR or AFR, at least in tubular organs, as we already proposed (20). In addition, the results correlate apical cell expansion with Crb subcellular localisation in the SAR. We propose that Crb accumulation in the SAR of LCJs promotes their growth facilitating the elongation of the cell along the longitudinal axis, in agreement with the proposed role of Crb promoting apical membrane growth (31, 32). Previous observations such as the expansion of the photoreceptor stalk membrane upon Crb overexpression (31) fully support this hypothesis, indicating that this can be a general mechanism.

## Conclusions

Here we find that during tracheal tube elongation Crb is transiently enriched in the SAR of DT cells in a planar polarised manner. This polarised distribution correlates with different dynamics or turnover of Crb protein, which appears to accumulate more at longitudinal junctions than at transverse ones. In addition, the polarised distribution correlates with the anisotropic growth of the apical membrane, axially-biased, that drives the longitudinal enlargement of the tracheal tubes. Our results are consistent with the notion that it is precisely the enrichment of Crb in the SAR of LCJs what promotes or facilitates polarised apical membrane growth in the longitudinal direction. Interestingly we also find that Src42A contributes to this planar polarisation of Crb, likely by promoting the preferential trafficking or recycling of Crb at the LCJs. Src42A was already known to regulate tube growth along the longitudinal axis, and we now propose that it performs this activity at least in part by promoting a Crb anisotropic enrichment. Src42A was also proposed to act as a mechanical sensor (17, 18, 29, 33–35), translating the polarised cylindrical mechanical tension (an inherent property of cylindrical structures) into polarised cell behaviour (17). Hence, we propose that Src42A would sense differential longitudinal/transverse tension stimuli and translate them into the cell by polarising Crb-mediated apical membrane growth in the longitudinal direction, thereby helping to orient cell elongation and as a consequence longitudinal tube growth.

In light of our results and previously published work we propose the following model (Fig. 5). Different mechanisms operate to regulate tube growth. On the one hand secretion drives apical membrane growth along the transverse axis independently of Src42A (17, 36). In addition, base level accumulation of Crb independent of Src42A (this work) may promote or contribute to isotropic apical expansion (16). On the other hand the presence of a properly organised luminal aECM also controls tube growth by restricting tube elongation (13, 14, 19). A Src42A-dependent mechanism acts in coordination with these other mechanims. Src42A would contribute to tube elongation through interactions with dDaam, the remodelling of AJs (17, 18) and topping up Crb accumulation at LCJs (this work). This increased accumulation of Crb at LCJs would bias the growth of the tube along the longitudinal axis, counteracting the restrictive activity of the aECM on tube elongation (13, 14, 19). In the absence of Src activity, the Src independent mechanism/s of membrane growth would still operate, and would favour a compensatory growth along the transverse axis as observed (17, 18), as diametrical growth is not restricted by the aECM. The regulation of size and shape of tubular organs is important for organ function, as evidenced by the fact that loss of regulation can lead to pathological conditions such as polycystic kidney disease (PKD), cerebral cavernous malformation (CCM) or hereditary hemorrhagic telangiectasia (HHT) (8, 37–39). Src proteins have been implicated in malformations like PKD (40, 41), highlighting the importance of investigating the mechanisms underlying their activities. While Src42A was proposed to regulate polarised cell shape changes during tracheal tube elongation through interactions with dDaam and the remodelling of AJs (17, 18), no polarised downstream effectors have been identified up to date. Hence, identifying that Crb anisotropy is one of the downstream effects of Src42A activity adds an important piece to the puzzle. Src42A and Crb are conserved proteins with important roles in development and homeostasis and are involved in different pathologies (41, 42). This work provides an ideal model where to investigate the molecular mechanisms underlying their activities, their interactions, and their roles in morphogenesis.

**Figure 5:**
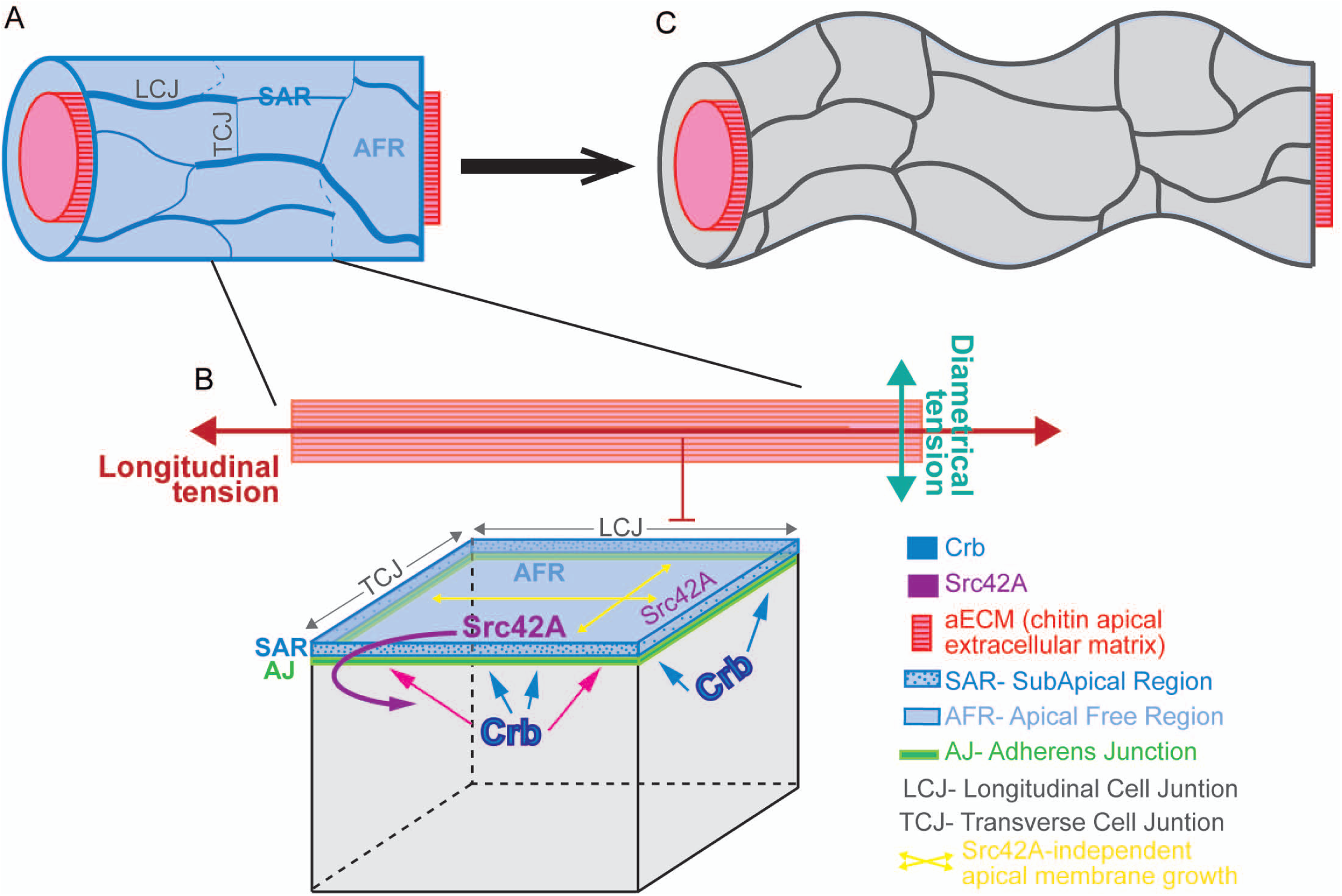
Model for DT elongation. Different mechanisms regulate the DT elongation. Crb protein accumulates apically in tracheal cells and becomes enriched in the SAR during tube elongation through a mechanism that is independent of Src42A (A and blue arrows in B). This base level of Crb may allow/promote an isotropic apical growth of tracheal cells (yellow arrows in B). Secretion also promotes a Src42A independent mechanism of apical membrane growth (yellow arrows in B). The aECM restricts the growth of the tracheal tube along the longitudinal axis (red arrow in B). Src42A, accumulated at cell junctions, senses the intrinsic differential tension along the tube axes and/or the aECM (B). Src42A promotes an increased accumulation of Crb at LCJs (pink arrows), resulting in Crb anisotropy (A). Crb enrichment in the SAR of LCJs, in turn, promotes apical membrane growth in the longitudinal direction, counteracting the restrictive activity of the aECM. The oriented cell elongation leads to the longitudinal tube growth (C).

## Materials and Methods

### Fly stocks

The following stocks are described in Flybase: *y ^1^w ^118^* (used as the wild type, WT, strain), *UAS-Egfr^DN^*, *UAS-Egfr^CA^* (*UAS-Egfr^λtop^*) (kindly provided by M. Freeman),UAS-Src42A^DN^ and UAS-Src42A^CA^ (kindly provided by S. Luschnig).

The *btlGal4* line (or *btlGal4-UAS-Src-GFP* to also mark the tracheal cells) was used to drive transgene expression in all tracheal cells from invagination onwards and *fkhGal4* to drive expression in the salivary glands.

The knock in allele *Crb^GFP^-C* was kindly provided by Y. Hong.

Control or embryos expressing the transgenes were collected after 13-17 hours at 29°C to obtain stage 16 embryos.

### Immunofluorescent stainings

Immunostainings were performed on embryos collected on agar plates fixed for 20 minutes (except for *D*E-cad staining, for which embryos were fixed for 10 minutes) in 4% formaldehyde in PBS. The following primary antibodies were used: mouse anti-Crb (Cq4) (1:20), rat anti-*D*E-cad (DCAD2) (1:100) from Developmental Studies Hybridoma Bank, DSHB; and rabbit anti-Serp (1:300) generously provided by S. Luschnig. Alexa Fluor 488, 555, 647 (Invitrogen) or Cy2-, Cy3-, Cy5-Conjugated secondary antibodies (Jackson ImmunoResearch) were used at 1:300 in PBT 0.5% BSA.

### Image acquisition

Fluorescence confocal images of fixed embryos were obtained with Leica TCS-SPE system using a 63x (1.40-0.60 oil) objective. Unless otherwise indicated, images shown are projections of Z stacks sections (0.2-0.4 μm).

Images were imported into Fiji (ImageJ 1.47n) and Photoshop for measurements and adjustments, and assembled into figures using Illustrator.

### Analysis of protein accumulation

Images of DT fragments between metameres 7 to 9 of stage 16 embryos were taken to analyse the Crb and *D*E-cad accumulation at cell junctions. After setting the longitudinal tube axis to 0°, cellular junctions were identified as longitudinal (LCJs, those oriented 0°±30° with respect to the axis) or transverse junctions (TCJs, oriented 90°±30°). Between 80-90% of junctions could be unequivocally assigned as LCJs or TCJs in all conditions analysed. The projection of the *D*E-cad channel was used to properly follow the cellular junctions to measure *D*E-cad and Crb protein accumulation. Accumulation of Crb or *D*E-cad was measured in all those junctions that were identified as LCJs or TCJs to avoid biased selection (i.e. in 80-90% of junctions of each metameres). To show the accumulation of proteins we generated a projection from the different stacks using the “Sum Fluorescence Intensity projection” tool in the Fiji software. To measure the total fluorescence at each cell junction we obtained the “raw intensity density” (the sum of all pixel values of the selected junction) manually drawing the junctions using the polygon selection tool of the Fiji software. The fluorescence of the proteins was normalised to the length of each junction. For each control or mutant embryo the ratio of intensity/length at LCJ and TCJ was calculated for Crb and *D*E-cad. The ratios were compared between control and Src42ADN embryos using the Scatter Plot tool of GraphPad Prism.

### FRAP assay

*btl*Gal4; *Crb^GFP^-C* (i.e. *control*) and *btlGal4-UASSrc42A^DN^*; *Crb^GFP^-C* (i.e. mutant) embryos were collected at 29°C. Embryos were mounted and lined up on Menzel-Gläser cover slips with oil 10-S Voltalef (VWR) and covered with a Teflon membrane (YSI membrane kit). FRAP was carried out on a Zeiss Lsm780 Confocal and Multiphoton System. The ZEN 2.1 SP3 of the Zeiss Confocal Software was used for data acquisition. The 488 nm emission line of an Argon laser was used for excitation at 1-2% power. We selected embryos at the appropriate stage (stage 15-16) oriented in a lateral position. We performed 10 pre-bleach scans with the 63x objective with a 1.5 zoom. These pre-bleach scans covered several Z-sections (1μm thick, 4 sections) in order to image the Crb accumulation at cell junctions that lie at different focal planes. Regions-of-interest (ROIs) of the same size were selected at longitudinal and transverse junctions. After 10 pre-bleach scans, bleaching was performed at these ROIs at high laser power (100 iterations at 100% power). Post-bleach scans were obtained immediately after bleaching at every 10 seconds and during 10-20 minutes. Average projections of the Z-sections were exported for each time point and assembled into a movie using Fiji software. The StackReg plugin in ImageJ was used to correct the movement of the embryo in the xy plane.

Fluorescence intensity in the ROIs was measured at each time point with Fiji software intensity plot profile tool. Igor Pro software was used for normalization, curve fitting (single-exponential fit), and calculation of recovery half-time (t_1/2_) and the mobile fraction (M_f_). The FRAP experiments were technically challenging. Although we performed FRAP for many (>20) ROIs from various embryos for each genotype, the trachea often moved during imaging, so many bleached ROIs were lost during recording. In addition, to FRAP ROIs at specific junctions in tracheal cells (with lengths typically ranging from 2,5 to 9 μm) was often difficult. Only movies where the ROIs remained in focus throughout (assessed by the fact that neighbouring junctions remained in focus) are reported here. Kymographs were generated using the dynamic reslice of the Fiji software.

### Crb subcellular accumulation

We quantified the total levels of Crb (using the Sum Fluorescence Intensity projection in Fiji) in different apical subcellular domains of embryos at stage 16. We selected individual cells from a region in the DT between metameres 7 and 9 or in the SG and generated projections of a few sections to include only the whole cell or a small number of them. We quantified Crb accumulation in the SAR by outlining the cell contour (using *D*E-Cad to visualise it) drawing a 6-pixel line on the junctional area and measuring the signal within each line. To measure Crb in AFR, a section inside the cell (defined by *D*E-cad junctional outline) was drawn with the freehand tool of Fiji. We expressed the subcellular accumulation as the ratio between SAR/AFR.

### Apical Surface Area measurements

We analysed the apical surface area of stage 16 tracheal and salivary gland cells, typically the same ones for which we analysed Crb accumulation in the SAR and the AFR. We selected individual cells from a region in the DT between metameres 7 and 9 or in the SG and generated projections of a few sections to include only the whole cell or a small number of them. We used maximal projections of *D*E-cad staining to outline the whole apical surface. We quantified the apical surface area by using the Freehand selection tool of the Fiji software, tracing a line following *D*E-cad staining. We expressed the apical surface area in μm^2^.

### Quantifications and Statistics

Total number of cells/embryos is provided in text and figures. Error bars indicate standard error (s.e.) or standard deviation (s.d) as indicated. *p*-values were obtained with an unpaired two-tailed Student’s *t*-test using STATA 12.1 software. \**p*<0.05, 0.001>\*\**p*<0.01, \*\*\**P*<0.001.

## Author contributions

M.L. designed the research and wrote the paper. I.O-C performed the experiments. I.O-C. and M.L. analysed and interpreted the data.

## Acknowledgements

We thank the contribution of M. Fuertes during the initial stages of this work. We thank E. Rebollo from the MIP-IBMB-PCB facility for imaging expertise. We are indebted to J. Colombelli from the ADM-IRB facility for help with FRAP experiments. We acknowledge the Developmental Studies Hybridoma Bank and the Bloomington Stock Centre for fly lines and antibodies. We thank S. Luschnig, M. Freeman and Y. Hong for reagents and flies. Thanks also go to the members of the Llimargas lab for helpful discussions and to M. Milán, J. Casanova, M. Furriols, S. Ricardo and A. Letizia for critically reading the manuscript.

This work was supported by Ministerio de Economía y Competitividad of the Spanish Government (BFU2012-39509-C02, BFU2015-68098-P) and from AGAUR of the Generalitat de Catalunya (2009 SGR1333). I.O-C. is supported by a FPI fellowship from Ministerio de Economía y Competitividad (BES-2013-065462). The funders had no role in study design, data collection and analysis, decision to publish, or preparation of the manuscript.

The authors declare no competing interests

---

**Figure S1:**
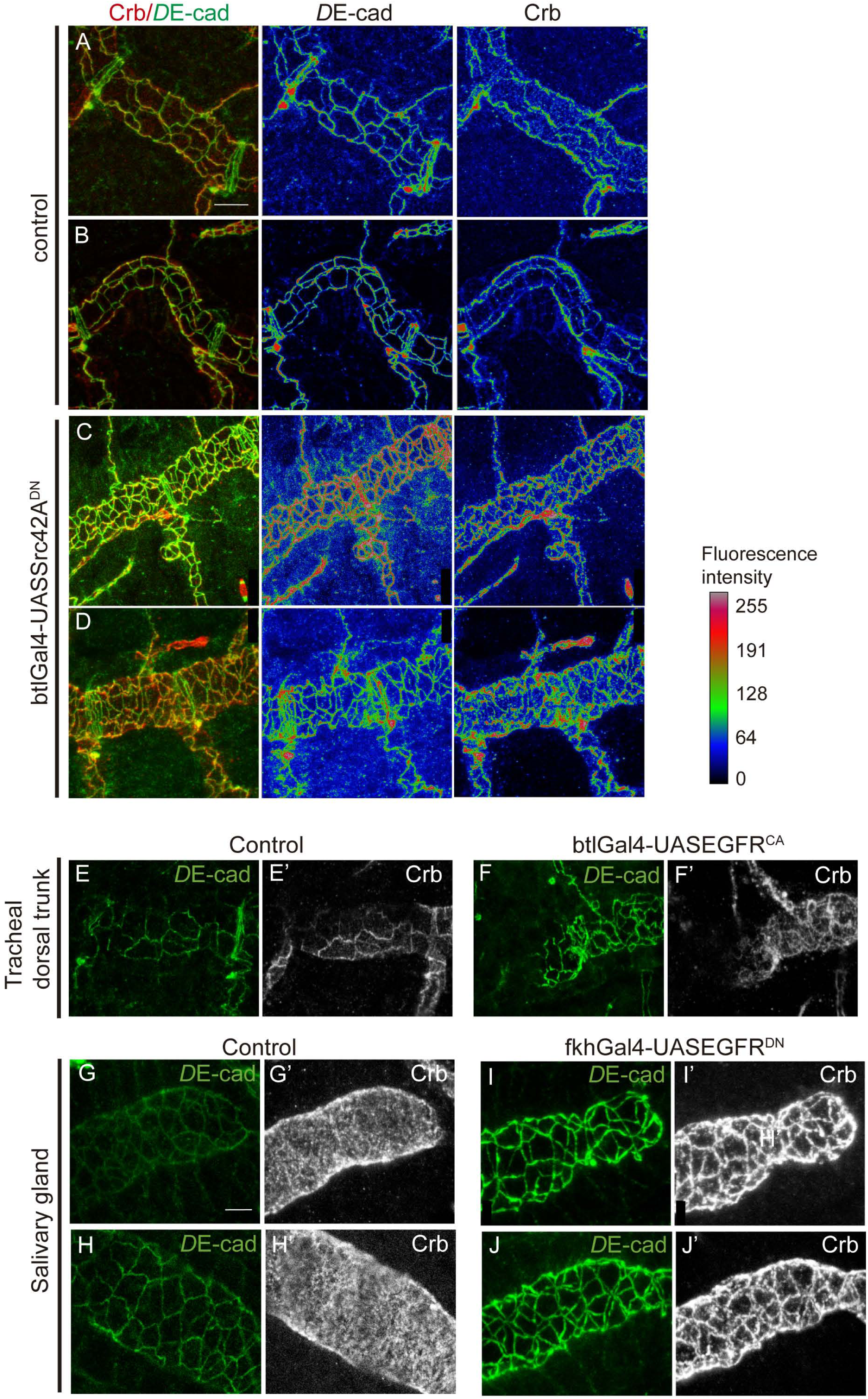
Accumulation of Crb and DE-cad in the DT and salivary glands. Further examples of Crb accumulation in control and different mutant conditions (A-D) Lateral views showing a region of the DT of stage 16 embryos of indicated genotypes stained for the indicated markers. The colour-coded fluorescence intensity shown for *D*E-cad and Crb matches the heat map shown on the right. Note that in contrast to the control (A,B), the fluorescence intensity in LCJs or TCJs for *D*E-cad and also for Crb is more homogeneous when Src42A is downregulated (C,D). Scale bar 10 μm (E-F) Lateral views showing a region of the DT of stage 16 embryos of indicated genotypes stained for the indicated markers. When EGFR is constitutively activated Crb accumulation in the SAR is less conspicuous and cells do not expand as in the control. Scale bar 7,5 μm (G-J) Lateral views showing a region of the salivary gland of stage 16 embryos of indicated genotypes stained for the indicated markers. Crb is localised in the whole apical area in control embryos (G’,H’) but when EGFR is downregulated it gets enriched in the SAR (I’,J’). G,I show projections of several confocal sections while H,J show a single confocal section. Scale bar 5 μm

**Figure S2:**
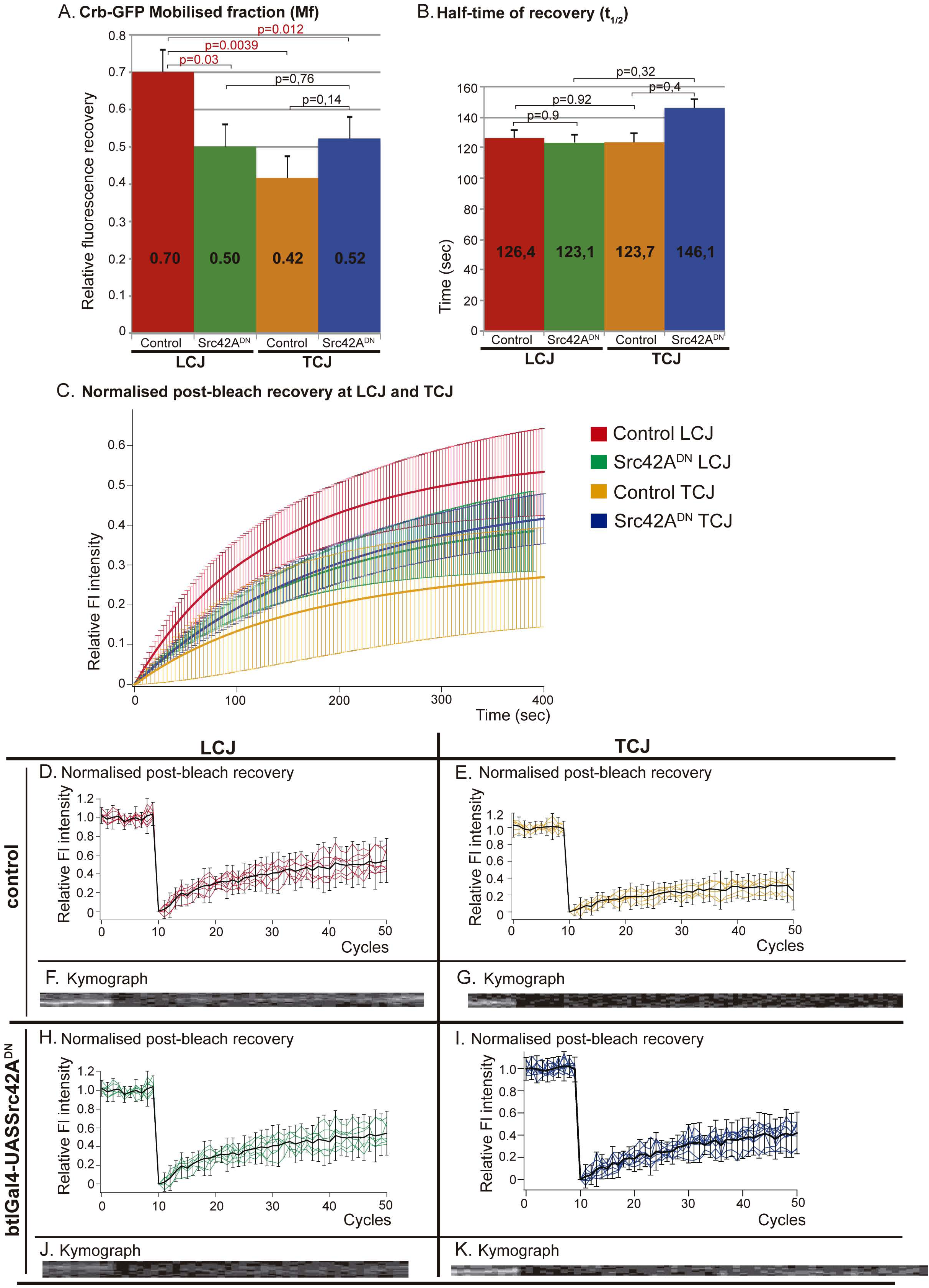
Analysis of crb-GFP FRAP experiments. (A) Comparison of the Mobile fraction observed at LCJs and TCJs in control embryos and in embryos in which Src42A is downregulated in the trachea. The Mf at control LCJs is significantly higher than in other conditions. Note that when Src42A is downregulated the Mf at LCJ decreases to levels similar to those of TCJs, suggesting that Src42A is required to increase the Mf precisely at the LCJs. (B) Comparison of the Half-time (t_1/2_), which is the time it takes to reach half of the Mf. The results indicate a similar kinetics of recovery in all conditions analysed. (C) Comparison of the fit of normalised post-bleach recovery of fluorescence at LCJs and TCJs in control and Src42A downregulation conditions. Note the higher recovery at LCJs of control embryos and the similar recovery at LCJs and TCJs in mutants and at control TCJs. (D,E,H,I) Normalised curves of FRAP experiments showing the levels of fluorescence relative to the pre-bleach levels during the cycles of image acquisition (a cycle every 10 sec). The samples were photobleached at cycle 10. The curves for each experiment (embryo) are shown in colours (red for control LCJs, orange for control TCJs, green for Src42^DN^ LCJs and bue for Src42^DN^ TCJs) and the average of the FRAP curves for each experimental condition is shown in black. (F,G,J,K) Kymographs of bleached areas for a representative example of each experimental condition. The homogeneous recovery of fluorescence in the bleached area indicates that it is not due to lateral diffusion. The X axis represents time and the Y axis represents distance. Error bars indicate standard error (s.e.). n=7 LCJs and n=6 TCJs from wild type embryos, and n=7 LCJs and n=9 TCJs from mutant embryos.

Movie 1: FRAP experiment in control LCJ

Region of a DT of a stage 16 *btlGal4; Crb^GFP^-C* embryo visualised from a lateral view using an inverted Zeiss Lsm780 confocal with 63x Oil objective and a 1,5 zoom. After several pre-bleach scans, a LCJ (marked with white arrow) was bleached. Post-bleach scans were taken immediately every 10 seconds for 10 minutes.

Movie 2: FRAP experiment in control TCJ

Region of a DT of a stage 16 *btlGal4; Crb^GFP^-C* embryo visualised from a lateral view using an inverted Zeiss Lsm780 confocal with 63x Oil objective and a 1,5 zoom. After several pre-bleach scans, a TCJ (marked with white arrow) was bleached. Post-bleach scans were taken immediately every 10 seconds for 14 minutes.

Movie 3: FRAP experiment in a LCJ in Src42A mutant embryos.

Region of a DT of a stage 16 *btlGal4-UASSrc42A^DN^; Crb^GFP^-C* embryo visualised from a lateral view using an inverted Zeiss Lsm780 confocal with 63x Oil objective and a 1,5 zoom. After several pre-bleach scans, a LCJ (marked with white arrow) was bleached. Post-bleach scans were taken immediately every 10 seconds for 10 minutes.

Movie 4: FRAP experiment in a TCJ in Src42A mutant embryos.

Region of a DT of a stage 16 *btlGal4-UASSrc42A^DN^; Crb^GFP^-C* embryo visualised from a lateral view using an inverted Zeiss Lsm780 confocal with 63x Oil objective and a 1,5 zoom. After several pre-bleach scans, a TCJ (marked with white arrow) was bleached. Post-bleach scans were taken immediately every 10 seconds for 12 minutes.

